# Harnessing the power of technical and natural variation in 116 yeast datasets to benchmark long read assembly pipelines

**DOI:** 10.1101/2022.03.17.484703

**Authors:** Noah Gettle, Brigida Gallone, Kevin J. Verstrepen, Rike Stelkens

## Abstract

With increases in throughput and reductions in cost, long read sequencing has become the standard for most genome assembly projects and has opened up new avenues for large-scale genomic research. While more amenable to assembly than short-read sequence data, long-read datasets tend to have higher error rates. To address this problem numerous tools have been developed to correct reads before assembly and polish assembled contigs. Although, numerous studies have been conducted to assess or benchmark these tools, few capture the real variance in long read sequence data that might affect tool performance much less full pipeline performance. To address these shortcomings, we compiled a dataset containing long-read sequences of 116 different strains of brewers’ yeast, *S. cerevisiae*, gathered largely from public databases and evaluated different assembly-related tools as well as their interactions. We found that pre-assembly short-read error correction of long reads combined with post-assembly short-read polishing provided the best assemblies. We also found that correction/polishing steps with uncorrected long reads often lead to degradation of assembly quality. Finally, we show which tools and pipelines work best with different types of input data.

## Main Text

The advent and development of long read or third-generation sequencing (TGS) has greatly expanded the possibilities of modern genomic research (De Coster et al. 2021; Hotaling et al. 2021). In contrast to short read or next-generation sequencing (NGS) technologies such as Illumina that typically generate reads < 500bp long, TGS platforms regularly produce reads between 10-30kb (Lin et al. 2021; Liao et al. 2019). Extended read lengths enable simpler and more rapid genome assembly (Koren et al. 2013; Liao et al. 2019; Koren and Phillippy 2015). This combined with decreasing costs and increasing throughput has made TGS technologies the new standard for *de novo* genome assembly, allowing researchers to produce highly contiguous genomes with less effort and fewer errors (Jung et al. 2019; Sohn and Nam 2018).

TGS technologies, however, come with unique costs that can make using long reads for assembly and downstream analyses challenging. Platforms from both of the commonly used long read sequencing companies, Oxford Nanopore Technology (ONT) and Pacific Biosciences (PacBio), produce sequences that can have relatively high error rates (5%-25% compared to <0.5% for Illumina) (Amarasinghe et al. 2020). Since using uncorrected assemblies can lead to significant errors in downstream genome analyses (Watson and Warr 2019), new algorithms and software packages are constantly developed and published to address these challenges (Amarasinghe et al. 2021). Many of these new programs aim to improve upon three steps in the assembly process: read error correction, assembly, and assembly polishing. The principles behind error correction and polishing steps are the same, in that they both use mapping information from read alignments to detect and fix erroneous base calls. For error correction, erroneous bases are corrected in the raw long reads prior to assembly while polishing corrects base errors in assembled contigs (Dohm et al. 2020).

The constant development of tools aimed at improving the outcomes of each step will ultimately result in improved assemblies but determining the optimal tools and pipelines to process any particular long read dataset can be a major challenge (Amarasinghe et al. 2020). While publications of tool comparisons or benchmarking analyses associated with long-read assembly are not uncommon all benchmarking analyses face tradeoffs that limit how comprehensive they are and thus the scope of their utility (Zhang et al. 2020; Cherukuri and Janga 2016; Dohm et al. 2020; Gavrielatos et al. 2021; Chen et al. 2020a; Sun et al. 2021; Wick and Holt 2019). For instance, many assessements of program performance focus on few input data sets that may represent a wide range of genetic variation (i.e. diverse organisms) but have limited technical variation. Although simulated data can address this problem to some degree, accurate simulations require accurate knowledge of the real variation in read quality researchers encounter. While there are some benchmarking analyses that do use larger sample sizes (N > 10), these are focused exclusively on prokaryotic genomes (Wick and Holt 2019; Chen et al. 2020b; De Maio et al. 2019; Chen et al. 2020a). Moreover, most benchmarking analyses avoid complications associated with testing programs from multiple assembly steps in concert despite the potential for interaction effects.

While tradeoffs are inherent to any benchmarking study (including this one), we attempted to address some of the limitations common to other analyses of long read assembly tools. To this end we capitalized on the popularity of brewers’ yeast, *Saccharomyces cerevisiae*, as a model for genomic research. By scouring public databases, we collected long read sequence datasets for 116 different *S. cerevisiae* strains. These data represent most of the different technological changes associated with ONT and PacBio platforms and range widely in terms of sequence quality. We ran these data through some of the most popular error correction, assembly, and polishing tools in different combinations to evaluate their performance and how tool effectiveness correlates with platform and overall read quality. As a service to the community, we also recorded and, to some degree examined, errors that arose while attempting to use these tools and have recorded the solutions to them to the extent we could.

We show that variation in input data naturally leads to difference in outcomes with respect to best tools and pipelines; however, using a large dataset we found certain programs and pipelines that perform best, in general. In particular, we found that use of both short-read pre-assembly error correction and post-assembly polishing produce the highest quality assemblies, but that long-read error correction and polishing does not generally provide significant gains and in some cases can reduce assembly quality. Further, we show a trade-off between contiguity and completeness/sequence quality between processed ONT-based and PacBio-based assemblies. Additionally, we found that non-kmer-based tools that use alignments to assess assembly quality (e.g. BUSCO and Reapr) may provide misleading results when used on low sequence quality (i.e. high error rate) assemblies. Although limited to a single species, we hope this study provides clarification on best practices to use when assembling eukaryotic genomes.

## Methods

### Sequence data and Quality Control

The sequence data used in this study was primarily collected from public databases (Table S1). These data represent 33 different projects and 116 different *S. cerevisiae* isolates published between 2013 and 2021 on all PacBio platforms excluding Sequel II (RS I, RS II, and Sequel I). Although specific details relating to ONT sequencing methods (e.g. basecaller) likely have bearing on assembly results (Lin et al. 2021), these data are not made consistently available alongside the public databases. For all strains, corresponding paired-end short-read Illumina data was also available.

Illumina reads were trimmed using TrimGalore (Krueger 2019). Improperly unsplit PacBio reads were corrected using pbclip (Kolmogorov 2019). ONT reads were used without processing. We generated input read summary statistics using Samtools and read data mapped to the *S. cerevisiae* reference assembly (R64-3-1_20210421) using BWA (Li 2013) in the case of short reads and Minimap2 (Li 2021) in the case of long reads (Table S2 and Table S3). Additional long read statistics were gathered using longQC (Fukasawa et al. 2020).

### Program selection and pipeline

The number of programs that could be assessed for each assembly step was limited by computational time and the number of data sets being evaluated. The selection of programs utilized in this study represents our attempt at balancing program representation in terms of age, popularity, and usability with resource constraints. Programs we tested but ultimately proved difficult to compile, install, or use on our shared server running CentOS are also listed in Table S3.

Error correction programs are designed to reduce sequence errors in long read data to facilitate more accurate downstream assembly. This is generally done either through the use of supplemental higher-accuracy short-read data or algorithms that evaluate self-alignments of the long read sequences. We evaluated two short-read-based programs, FMLCR (Wang et al. 2018) and Ratatosk (Holley et al. 2021), and three long-read-based programs, Canu (Koren et al. 2017), MECAT2 (Xiao et al. 2017) (PacBio only), and NECAT (Chen et al. 2021) (ONT only). The resulting corrected sequences in addition to the raw reads were directly assembled using Canu, Flye (Kolmogorov et al. 2019), wtdbg2 (Ruan and Li 2020), or Shasta (Shafin et al. 2020). As Shasta was created primarily to assemble ONT-generated reads, we adjusted its parameters when assembling PacBio reads (consensusCaller = Modal; minReadLength = 5000). Otherwise, all error correction and assembly programs were run with the default settings specified for the corresponding long read sequencing platform.

Polishing, much like error correction, involves using long and/or short reads to improve sequence accuracy, although in an assembled genome rather than the input reads. To test the effects of the polishing step and the various programs designed to do it, we polished the highest-ranked assemblies for each input data set using Pilon (Walker et al. 2014), POLCA (Zimin and Salzberg 2019), NextPolish (Hu et al. 2020), ntEdit (Warren et al. 2019), Apollo, and Racon (Vaser et al. 2017). For Apollo and Racon, uncorrected long reads were used as input. Additionally, we tested the performance of Racon using FMLRC error-corrected reads as input. To determine the relative impacts of pre-assembly error-correction on downstream polishing, we also assessed polishing effects on the highest-ranked assemblies generated from uncorrected reads.

### Program errors

Not all combinations of input data and program successfully produced assemblies. For each program error we recorded or summarized the error and attempted to solve the error if, based on our judgement, it would not bias the outcomes of our assessments (Table S5). As most of these errors were generated by Canu, we looked specifically at the covariates that might affect successful assembly with this program. To do this we used a logistic generalized linear model testing the relative effects of all raw long read variables generated by Samtools’ samstats and longQC. We reduced the model using an AIC-optimized stepwise function followed by removal of covariates with variable inflation factors (VIF) greater than 4.

### Assembly assessment

Assembly quality can be broken down into four metrics: contiguity, completeness, assembly correctness, and sequence correctness. Contiguity, or the number and size of assembled contigs, was estimated using e-size (Salzberg et al. 2012). Completeness, or fraction of the true genome represented by an assembly, was estimated using the short-read kmer-based ‘completeness’ metric generated by Merqury (Rhie et al. 2020). Assembly correctness was measured in terms of misassemblies per contig as identified with REAPR (Hunt et al. 2013) while sequence correctness, referring to per base accuracy, was estimated using Merqury-generated QV values. We tested Spearman’s correlations between these metrics and other traditional descriptive assembly statistics such as N50s and BUSCO scores for all assemblies following error correction and assembly steps (Gurevich et al. 2013; Thrash et al. 2020). To determine how best to gauge assembly quality when supplemental short read data are unavailable, we also ran all assemblies through Quast (Reference assembly accession = GCA_000146045.2) (Gurevich et al. 2013).

Overall effects of error correction methods, assembly tools, and the interactions between them were tested separately for ONT-based (N = 48) and PacBio-based (N = 27) assemblies using a MANOVA with our four chosen quality metrics (e-size, completeness, misassemblies per contig, and QV) as response variables. Post-hoc Tukey HSD tests of factor comparisons were based on ANOVAs run independently for each response and FDR-adjusted to account for correlation in response variables. As there was a bias in the tools that produced errors rather than attempt assembly from reads of some datasets (Tables S5, S6), we only tested effects on datasets for which all error correction/assembly tool combinations resulted in assemblies.

We tested polishing tool performance and whether it differed based on whether assemblies were generated from corrected or uncorrected reads using a MANOVA with the difference in pre- and post-polishing metrics used as response variables. Polishing effects were tested using a Wald’s test of estimated marginal means. Post-hoc Tukey contrasts were performed on estimated marginal means with Bonferroni correction.

In order to rank assembly outputs from each program or set of programs, quality metrics were standardized using Z-scores (determined with respect to each input read set). The sum of these unweighted Z-scores was used to rank assembly outputs for each input read set following error-correction/assembly and polishing steps.

## Results

### Core Assembly Quality Metrics

Assessment metrics are often divided into those that serve as estimates of contiguity, completeness, and assembly accuracy (misassembly rates). Because long read data generally have high error rates, TGS-based assemblies can also be evaluated based on sequence accuracy (sequence error rate). To represent these metrics, we chose e-size, completeness (Merqury), number of missassemblies per kb (REAPR), and QV (Merqury). While significantly correlated, we found that the strength of these correlations was less than those between other metrics commonly used to describe similar aspects of genome assembly (e.g., N50 and e-size) (Fig. S1).

Outside of e-size, the main metrics we used require supplemental short-reads to generate. Although available for this study, we wished to determine the best way to assess assembly quality when short reads are not available. As such, we examined correlations between Quast generated statistics and our three core short-read dependent metrics. In terms of completeness, the total percent of identified BUSCOs (complete and partial) corresponded well with Merqury’s completeness metric (Fig. S2A; Spearman correlation; ρ = 0.91, p < 2.2e-16). The relationship between BUSCO scores and completeness was linear for assemblies generated with PacBio reads regardless of whether they were corrected or uncorrected. This relationship for short-read corrected ONT-based assemblies, however, was logarithmic (Fig. S2B). Additionally, there is an inverse logarithmic relationship between the frequency of indels detected by QUAST and QV scores (FigS2C). A combination of total number of BUSCOs, indel frequency, and e-size metrics predicted the highest ranking assemblies (as determined by QV, misassembly rates, correctness, and e-size) 65.5% of the time, significantly higher than random (one-sided exact binomial test, N = 116; p < 2.22e-16). The top assemblies, as predicted by these short-read-independent metrics, appeared in the three top-ranked assemblies based on short-read metrics 94.8% of the time.

### Error correction and assembly program performance

For each input data set, we generated assemblies using all combinations of the four tested error correction tools (Canu, MECAT2/NECAT, Ratatosk, and FMLRC) and assembly programs (Shasta, Canu, Flye, and wtdbg2). Additionally, we ran uncorrected reads through each assembler. For a number of read sets (37.9%), certain error correction–assembler combinations did not produce assemblies. In terms of complete failures (i.e., no assembly generated), Canu’s assembler was the most frequent culprit (36.2% of input reads failed in some combination), particularly in combination with FMLRC and Canu error correction (Table S6). Canu errors were generated due to reported insufficient coverage likely a result of strong read filtering by the error correction tool, the assembly tool, or both. Input datasets that failed Canu at least once were significantly associated with lower coverage (coverage mode) and higher error rates (mismatches per base mapped to the reference) in the raw reads (Logistic GLM; Fig. S3).

The relative impact of each error correction and assembly tool on genome quality varied considerably based on the sequencing platform (Fig. S4). For ONT datasets, short-read error correction resulted in better assemblies than uncorrected or self-corrected reads with regards to all metrics with the exception of contiguity (e-size). There were no significant differences between the two short-read error correction programs tested here (FMLRC and Ratatosk). Self-correction had no significant effect on final ONT-based assembly quality except when done in combination with Shasta assembly (Fig. 1). For PacBio-based assemblies, both types of error-correction significantly improved QV values in final assemblies while all error-correction tools, except for FMLRC, reduced misassembly rates. Otherwise, the impacts of error correction alone were largely tied to interaction effects with the different assembly tools (Fig. 2). With regard to assembly programs, Canu, Flye and wtdbg2 produced assemblies with similar QVs, misassembly rates, and levels of completeness when using ONT data. In terms of contiguity, however, wtdbg2 and Flye outperformed Canu. For PacBio reads, Canu and Flye consistently outperformed both Shasta and wtdbg2 across all quality metrics with Flye edging out Canu in terms of contiguity (e-size).

**Figure 1.**
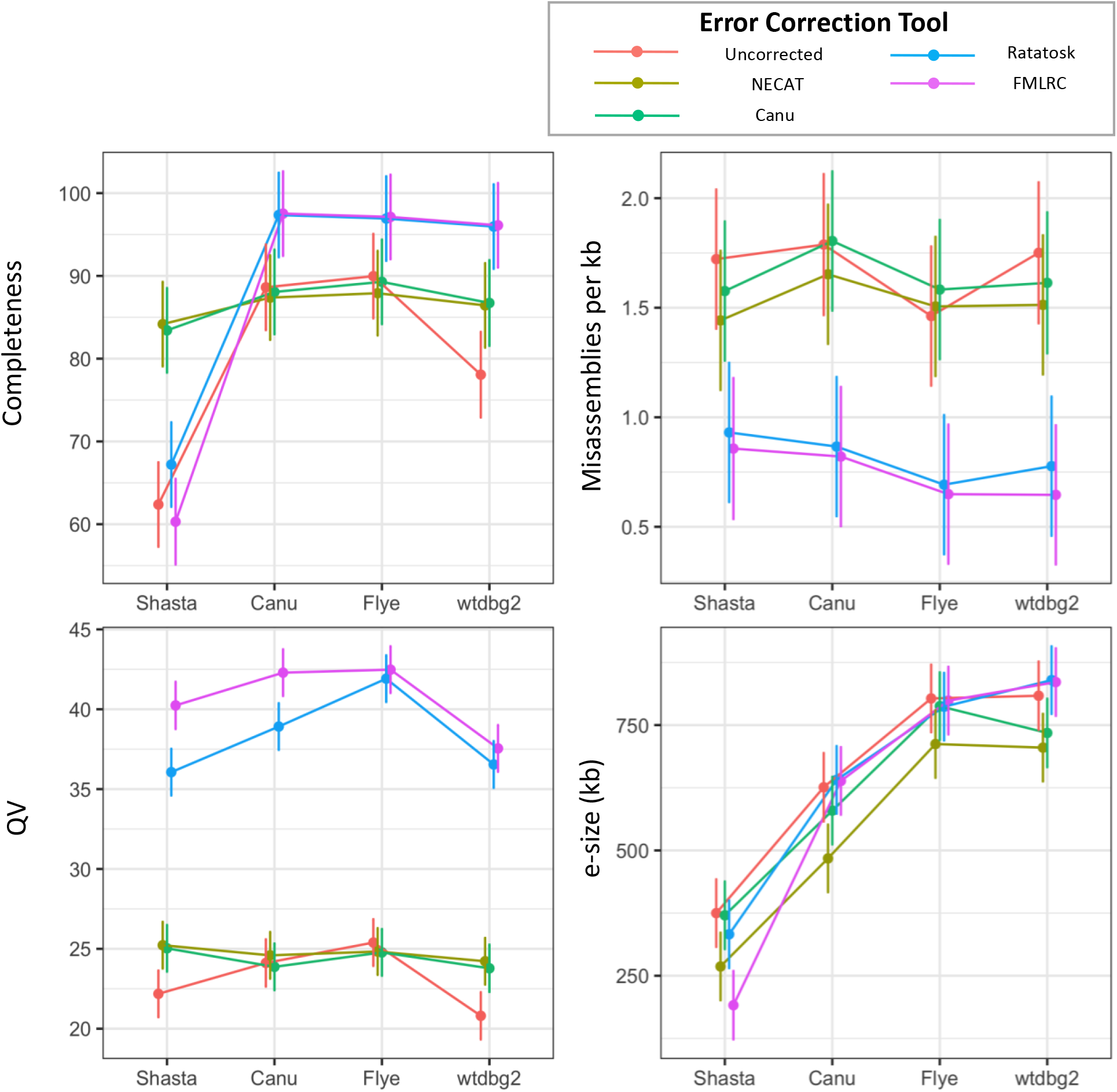
Effects of various error correction and assembly tools on ONT assembly quality. Estimated marginal mean values of four measures of assembly qualities for ONT read sets assembled using different combinations of error correction and assembly tools. Error bars = 95% CI.

**Figure 2.**
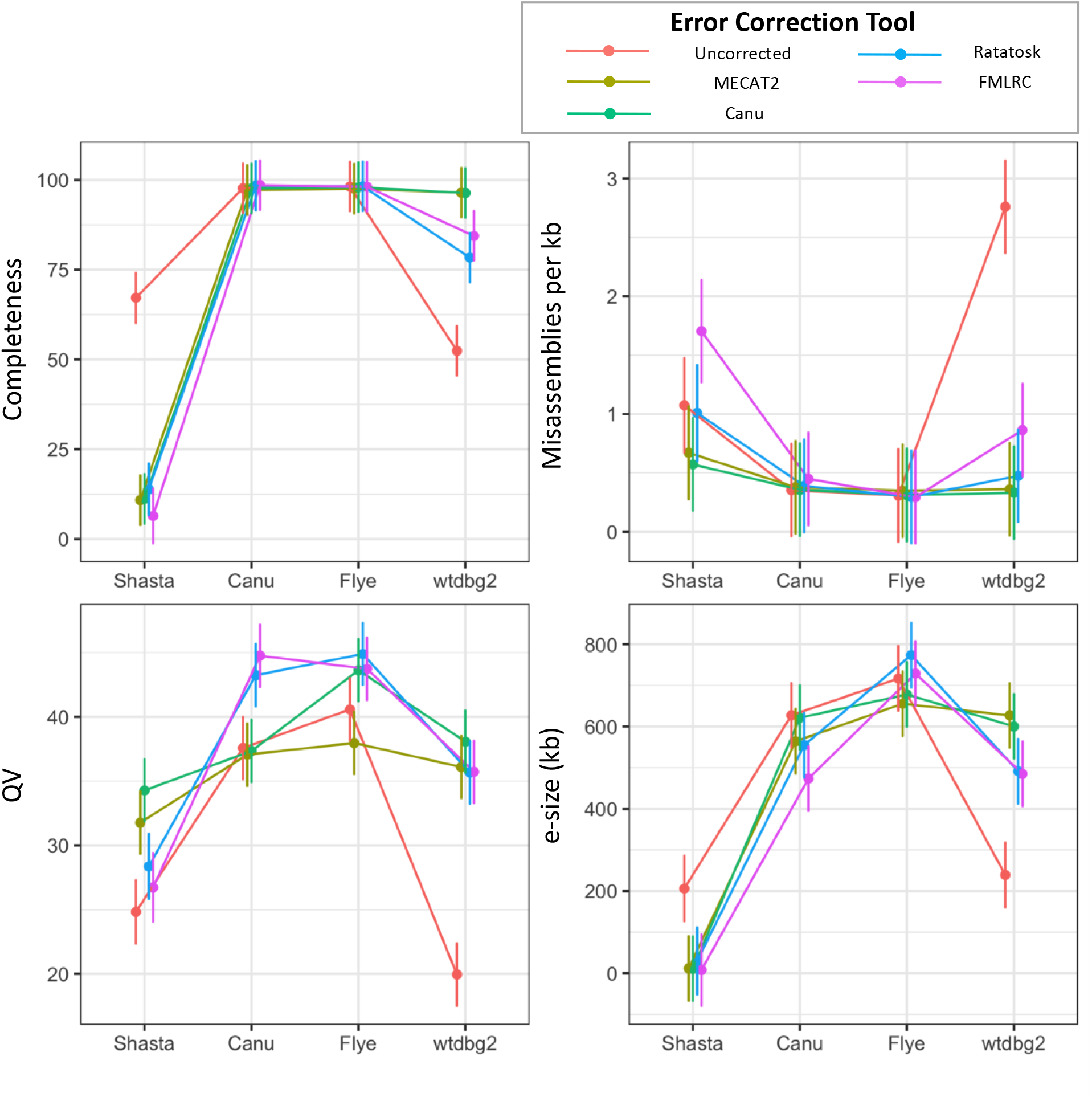
Effects of various error correction and assembly tools on PacBio assembly quality. Estimated marginal mean values of four measures of assembly qualities for PacBio read sets assembled using different combinations of error correction and assembly tools. Error bars = 95% CI.

In addition to testing the relative tool effects, we also ranked assemblies for each input dataset based on a combined unweighted estimate of overall quality. Assemblies deemed best based on our combined unweighted metric were not always or even generally optimal with regards to all metrics; however, we felt that this metric balanced tradeoffs well enough for use in selecting assemblies for further analysis (Fig. S5). Overall, Flye produced the greatest fraction of top ranked assemblies for both ONT (64%) and PacBio (56%) reads according to our combined quality metric (Fig. 4A). Among assemblies produced without supplemental short reads, Flye again proved to be most frequently the best all-around assembler (ONT = 69%; PacBio = 95%). Surprisingly, uncorrected reads more frequently (62%) produced better assemblies than those that were self-corrected (Fig. 4B).

**Figure 3.**
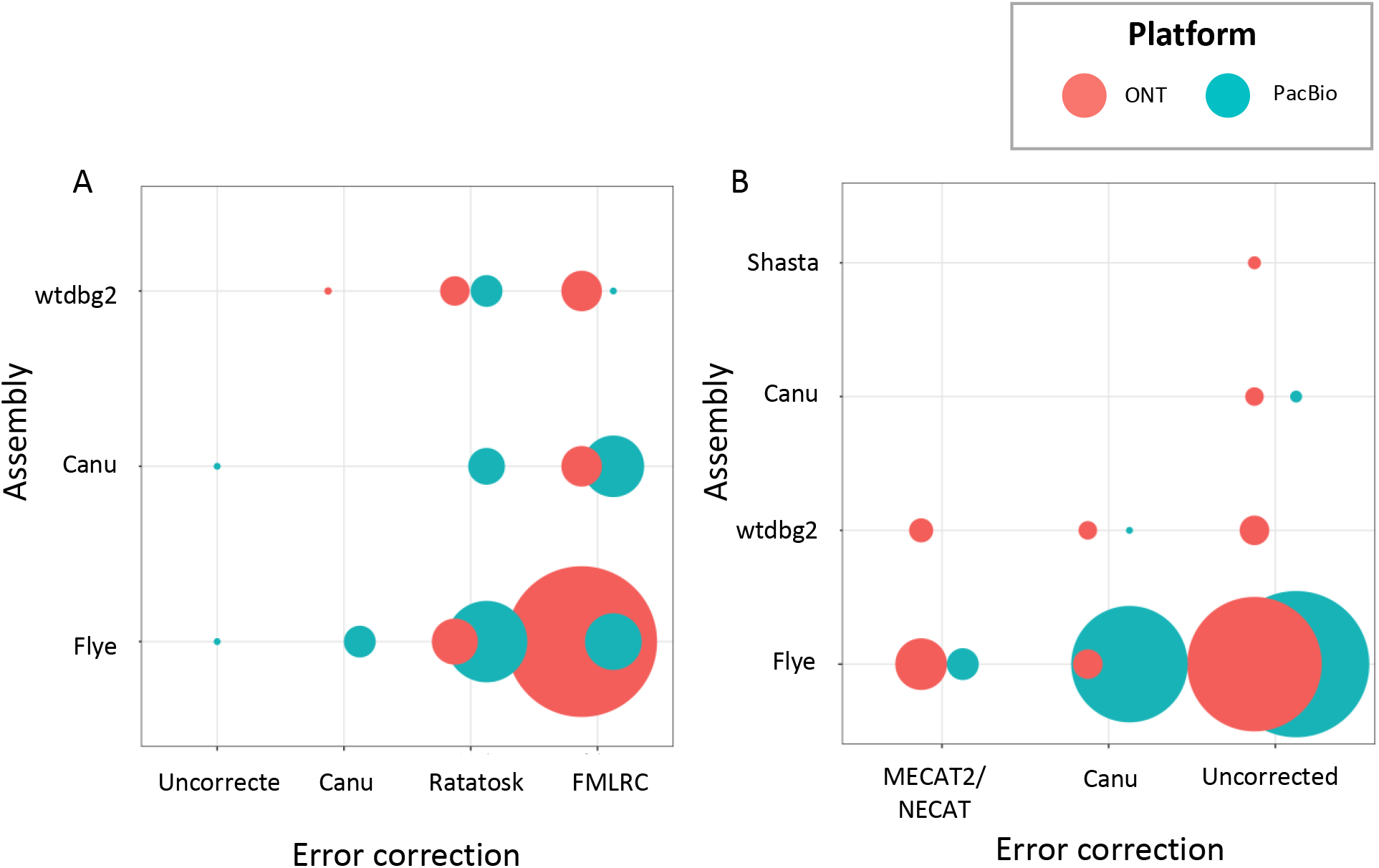
Flye produces the greatest number of highest-ranked assemblies. Fraction (corresponding to circle area) of assemblies ranked highest for each error correction / assembly tool combination. A) Overall top ranked assemblies B) Top ranked assemblies for error correction methods not requiring supplemental long reads.

**Figure 4.**
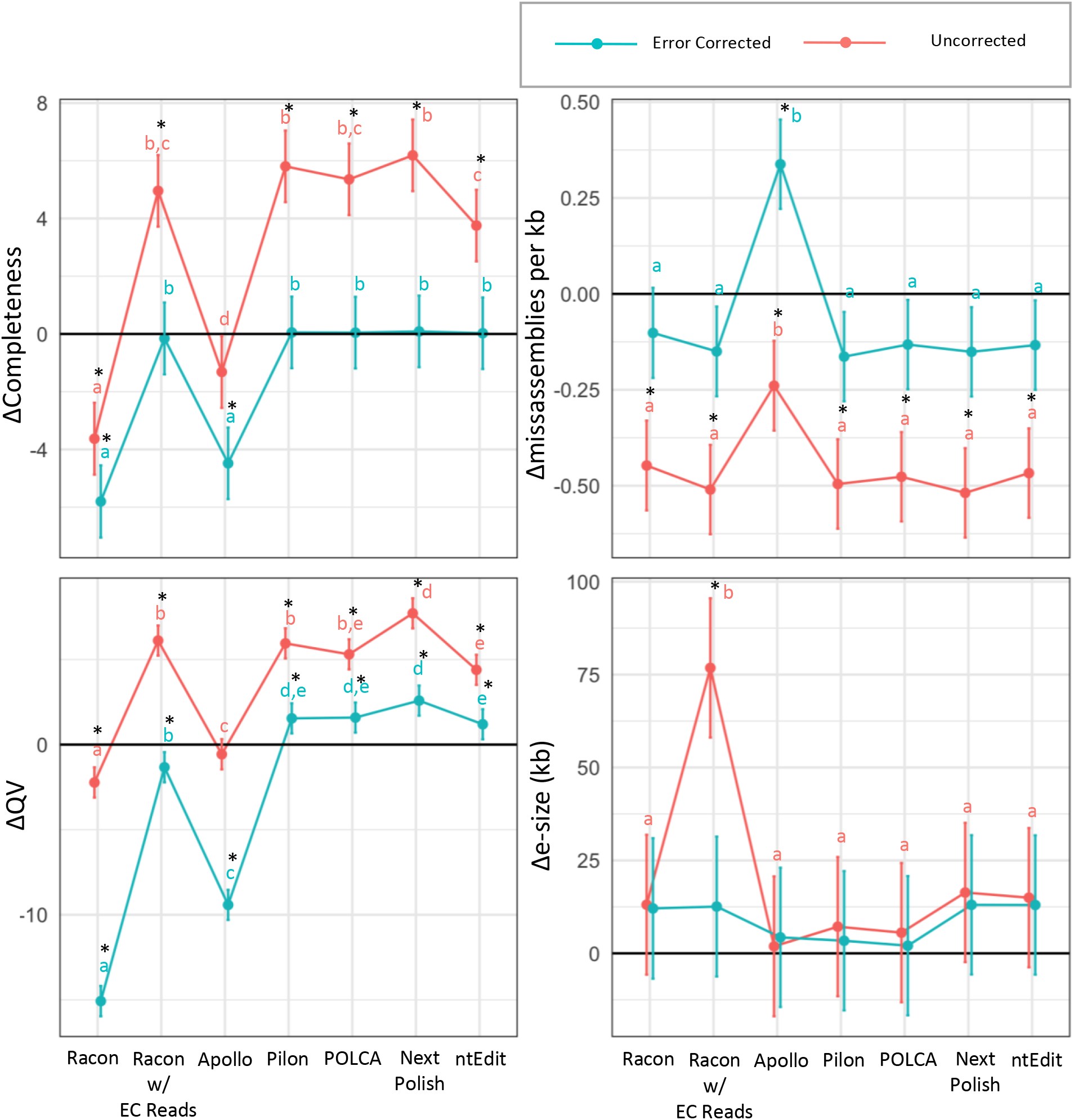
Effects of polishing on assembly quality. Estimated marginal mean change in values of four measures of assembly quality relative to input. Error bars = 95% CI. * = Value is significantly greater or less than 0 (t-test; Bonferroni adjusted p < 0.05). Letters = Significantly different groups (TukeyHSD; Bonferroni adjusted p < 0.05). Tests based on estimated marginal means derived from multivariate model. EC = FMLRC error corrected reads.

### Polishing tool performance

A common post-assembly processing step is to polish contigs using either supplemental Illumina short reads or long reads. In many ways, this step is analogous to pre-assembly read correction. There are no examples that we have found, however, that directly examine whether it is most effective to do pre-assembly read correction, post-assembly contig polishing, or both. To determine the relative effects of sequence polishing on assembly quality, we polished all of the top-ranked assemblies from our error correction/assembly step (primarily error-corrected) as well as the top-ranked uncorrected-read-based assemblies using six different tools. Four of these (Pilon, POLCA, NextPolish, and ntEdit) use short reads to correct sequence errors while one (Racon) uses long reads, and one (Apollo) uses a combination. With Racon, we additionally looked at the difference between self-polishing using uncorrected and FMLRC corrected long reads.

Short-read based polishing on error-corrected assemblies from the previous step resulted in far fewer significant gains in quality than correction of uncorrected assemblies across all metrics except e-size. Naturally, this might be expected given the lower input quality of uncorrected assemblies; however, the correlation between input quality and quality gains was not consistently linear and/or positive or negative across the different polishing tools and the different quality metrics (Fig. S6). Regardless, short-read polishing and Racon polishing with FMLRC long reads resulted in significant quality gains in terms of QV for both error-corrected and uncorrected assemblies with the addition of gains in completeness and misassembly rates for the latter (Fig. 4). Interestingly, polishing with uncorrected long reads (Racon and Apollo) resulted in average reductions in completeness and QV, particularly in error corrected assemblies. Relatively speaking, however, we found few significant differences between short-read-based polishing tools with the exception of major average gains in e-size among assemblies polished using Racon and FMLRC error-corrected reads. Taken as whole, however, we found that polishing with NextPolish overwhelmingly tended to result in the greatest gains in the highest fraction of input datasets except in the case of uncorrected-PacBio-read-based assemblies (Fig. S7).

A common tactic to improve assembly quality is to polish assemblies multiple times, typically with the same program or with a combination of long-read and short-read polishing programs. We examined the effectiveness of this technique by subjecting our top-ranked error-corrected assemblies to 10 rounds of polishing with our highest ranked short-read polisher (Fig. S8). More than 1 round of polishing did not significantly improve either ONT or PacBio assemblies, although most assemblies gained some small amount of completeness after the second round. Additionally, we looked at the effects of polishing once with Racon using FMLRC-error-corrected long reads followed by multiple rounds of polishing (Fig. S9). While Racon polishing resulted in significant increases in contiguity in both PacBio and ONT assemblies, PacBio assemblies on average also suffered significant reductions in QV and completeness. One round of short read polishing was largely able to counterbalance the effects on QV but not on completeness.

### Error correction, polishing, or both?

While error correction and polishing are both, in theory, quite similar, rarely are their effects compared with each other. As such it is unclear whether it is preferable to do both steps or if there are interaction effects that might make use of one or the other preferable. To compare error correction and polishing steps we examined estimated marginal mean quality values for the top assemblies for each sample produced with and without error correction and with and without polishing (Fig 5).

**Figure 5.**
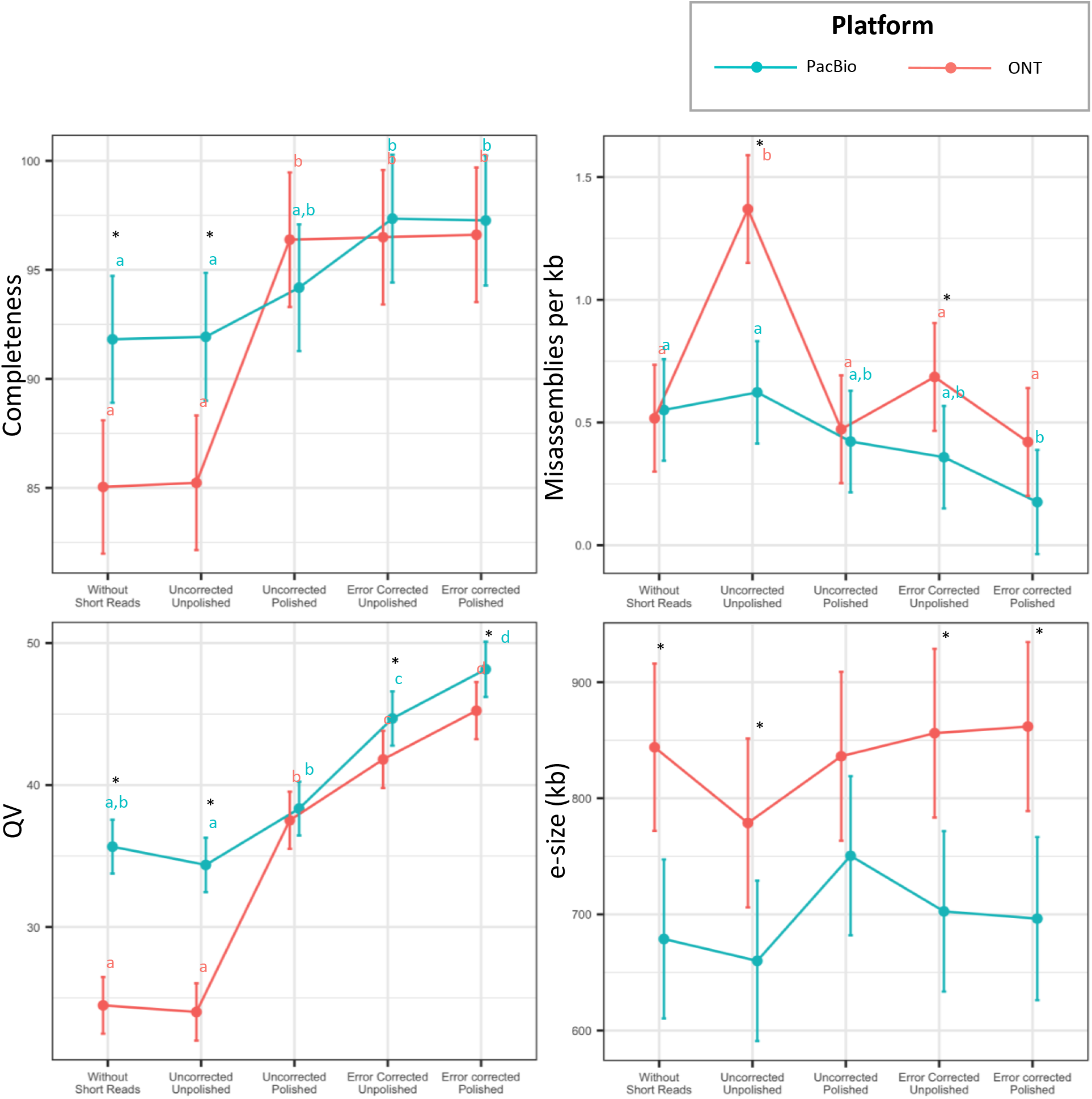
Effects of short read processing, error correction, and polishing on assembly quality. Estimated marginal means for quality estimates of top assemblies from each assembly category. Error bars = 95% CI. * = Values for different platforms significantly different; Letters = Significantly different groups (TukeyHSD; Bonferroni adjusted p < 0.05). Tests based on estimated marginal means derived from multivariate model.

Short-read assisted methods (error correction, polishing, or both) tended to improve assemblies with regards to all quality metrics outside of e-size (contiguity). Interestingly, preassembly error correction had a greater impact on QV than polishing although assemblies having undergone both steps had the best mean completeness, missassembly, and QV scores. With regards to platform, error correction and polishing significantly reduced quality differences between ONT- and PacBio-based assemblies; however, ONT consistently had higher levels of contiguity (e-size) while PacBio assemblies had greater levels of sequence quality (QV).

### Best assembly pipelines

In addition to comparing the different steps in the assembly process and programs available for each step, we wished to determine which pipelines generated the best assemblies. Breaking down programs based on the frequency in which they appear in the top pipeline for a given input data set, we found that the assembly program Flye and the polisher NextPolish were involved in producing a large majority of the highest quality assemblies (58.6% and 70.0% respectively) compared to the next best assembler (Canu = 24.1%) and polisher (Racon w/ FMLRC-corrected reads = 14.7%) (Fig. 6). For error correction, the best tool for ONT error correction was clearly FMLRC (69.1%) whereas Ratatosk and FMLRC performed similarly for PacBio error correction (47.3% and 39.3% respectively) (Fig 6). When just considering assemblies produced without additional short reads, uncorrected reads supplied to Flye produced the best results. While Racon largely seems to have improved ONT assemblies, post-assembly polishing with long reads seems to largely reduce quality in PacBio assemblies (Fig. S10)

**Figure 6.**
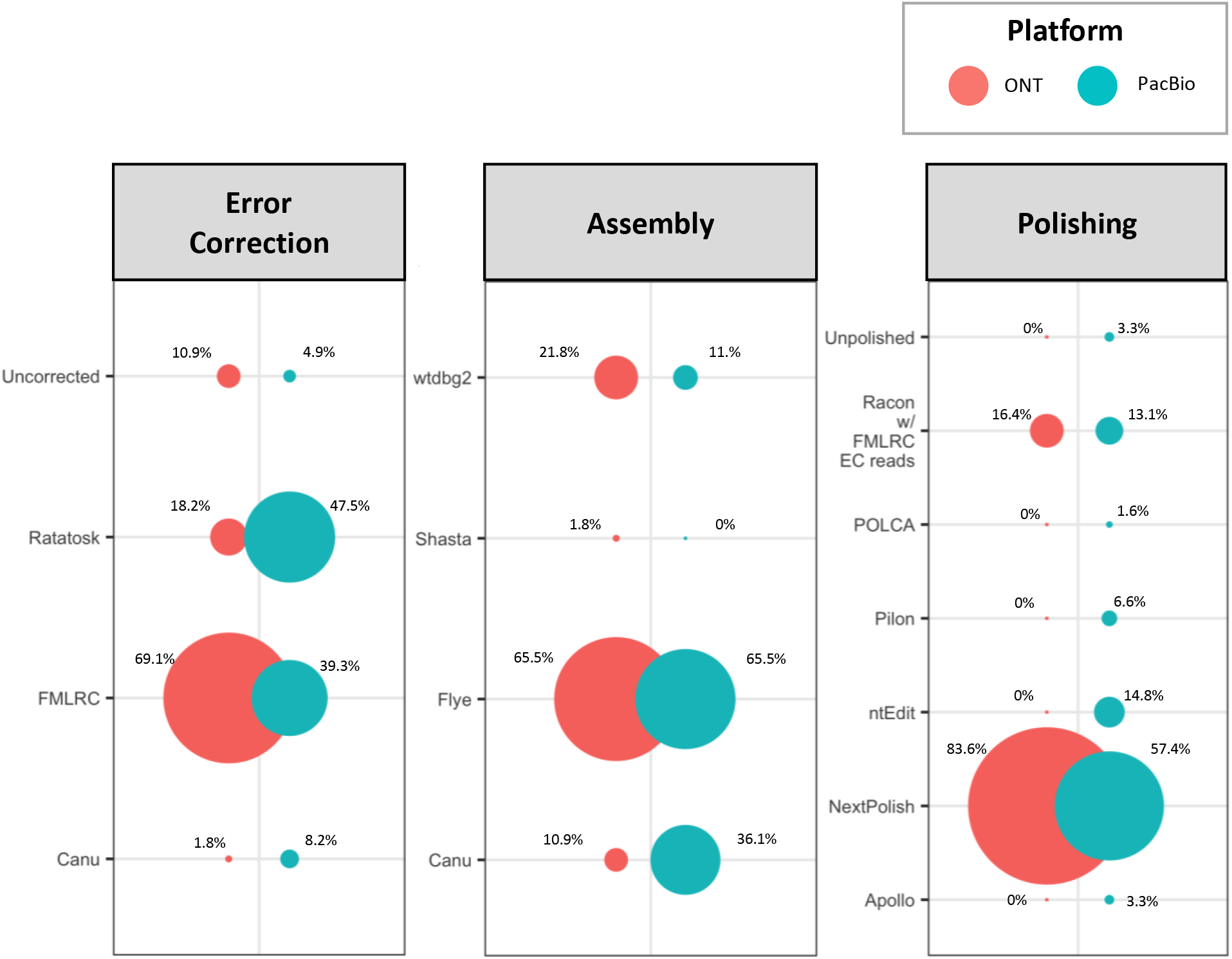
Top assembly pipelines. Fraction (corresponding to circle area) of assemblies ranked highest for each error correction, assembly, and polishing tool combination.

## Discussion

### Assembly assessment

There are four aspects of a genome assembly that one should consider when assessing its quality: contiguity, sequence accuracy, assembly accuracy, and completeness. Unless a closely related reference genome is available, one has to make use of proxy metrics that may not entirely be reflective of the aspect of assembly quality we wish to measure. One way to avoid such issues may be to use external data such as short-read kmer mapping to assess assembly quality as suggested by Rhie et al. (2020). In this study, we used the metrics Rhie et al. suggest as well as misassembly rates as measured using the tool Reapr to not only evaluate tools and rank assemblies but also to see how they correlate with other common measures of assembly quality.

As might be expected, in most sets of assemblies we found a strong positive linear correlation between the fraction of BUSCOs identified and Merqury’s completeness metric (Fig S2). This was not, however, true in ONT-based assemblies that had not been corrected with short-reads where, although the correlation was positive it followed a logarithmic pattern. This is rather concerning and indicates that even when a high-quality BUSCO set is available, in some conditions, this metric might not accurately reflect completeness. Indeed, closer examination shows that the BUSCO score metric is more sensitive to high error rates in the input reads (error rate > 20%) which may reflect alignment issues and thus problems with sequence accuracy rather than completeness (Fig. S11).

Using Reapr to measure misassembly rates, we also encountered potential issues. Although clearly correlated with our measured sequence accuracy (i.e. Merqury QV), we felt like the program might provide a useful reference-free way to measure assembly accuracy. While this might be the case, it was clear after seeing significant effects of polishing (which should not alter assembly arrangement) on misassembly rates that correlate with changes in QV that Reapr’s detection of misassemblies and, thus, its measure of sequence accuracy may, reflect poor sequence accuracy as well (Fig S12). Together, these data suggest that one should be careful when using alignment-based metrics such as BUSCO scores or Reapr-detected misassembly points to assess assembly quality when input long reads have high error rates that have not been corrected via supplemental low error rate short reads.

### Best Practices

In most cases we found that it was best to both correct reads before assembly with short reads and either FMLRC or Ratatosk as well as to short-read polish after assembly. For the polishing step, we found that NextPolish tended to produce the best assemblies. However, Racon polishing with FMLRC-corrected long reads resulted in significant improvements in contiguity suggesting that, in cases where increasing contiguity is the goal, long-read polishing with error corrected reads might produce desired gains. For PacBio-based assemblies, however, it should be noted that long read polishing can come at a cost to other quality metrics that may not obviated via subsequent rounds of short-read polishing. For ONT-based assemblies, the optimal pipeline appears to be one round of error-corrected long read polishing followed by one round of short read polishing.

In the case where supplemental short reads are not available, our results suggest that it is generally best not to error correct reads and for, PacBio assemblies, to avoid polishing. For Nanopore assemblies, polishing with Racon may result in some improvements. Additionally, our results suggest that more than 1 or 2 rounds of polishing with the same tool will likely not result in significant gains in quality.

While both Canu and Flye assembly algorithms were capable of producing high quality genomes, Flye tended to produce the best assemblies most frequently according to our combined metric. The primary difference between Canu and Flye came down largely to output assembly contiguity. Moreover, Canu is much more sensitive to input data coverage/quality than the other tools tested here and, without advanced settings adjustments, frequently errors out.

Naturally, the given starting point for any assembly project is choosing the appropriate sequencing platform. Based on this study, it is clear that supplemental short reads go a long way towards improving assembly quality, especially when working with ONT long reads. Still with our three-step assembly process ONT-based assemblies were on average of lower quality except when considering contiguity.

### Conclusion

While we cannot claim that our study is comprehensive with regard to the range of diversity in organisms and tools we assessed, our approach, using a greater number of comparable input data sets, provides new insight into how assembly tools function in relation to variation in input data and other long-read assembly tools. We were also able to statistically assess which pipelines and tools generally work best for ONT- and PacBio-based data and for long-read assemblies where supplemental short reads may or may not be available. There was, however, considerable variance in outcomes of tools that we were unable to correlate with any input data measurements (e.g. read error rate). While some of this may be a result of ‘noise’ (e.g. thread seeding in Flye), there may be more input data characteristics that determine which program or pipeline works best. Additionally, there are numerous tools and assembly approaches we were unable to evaluate here. We think, however, that future approaches akin to ours may help reveal even better approaches to long read genome assembly.

## Acknowledgements

We thank Meredith S. Palmer and the members of the Stelkens lab for statistical and conceptual advice.

## Author contributions

Conceptualization: NG

Data curation: NG

Formal Analysis: NG

Funding acquisition: RS

Investigation: NG

Methodology: NG

Project administration: NG, RS

Visualization: NG

Writing: NG, RS

## Data and materials availability

Code and data are available upon request.

