## Supplementary Figures for "Harnessing the power of technical and natural variation in 116 yeast datasets to benchmark long read assembly pipelines"

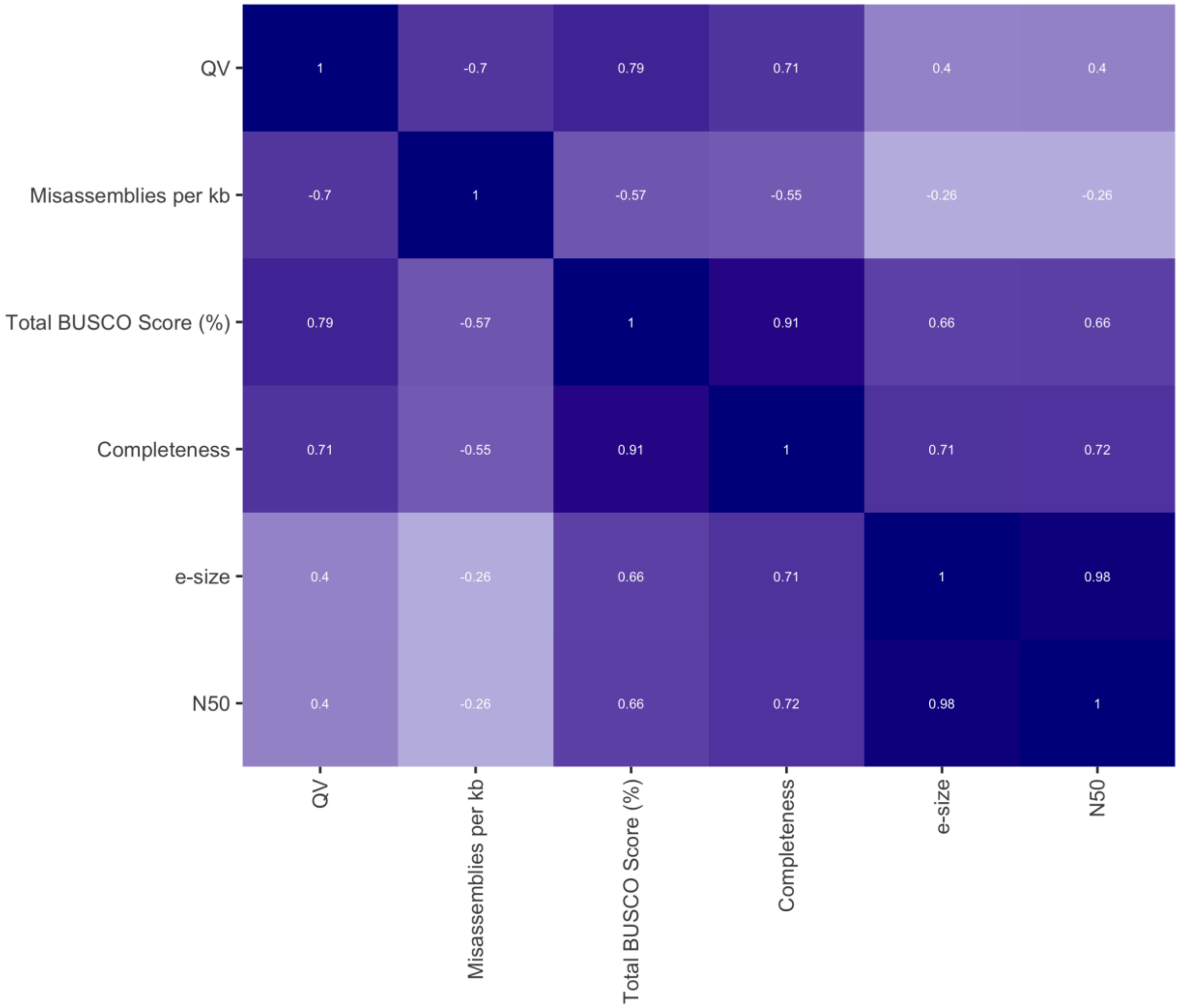


**Figure S1. Correlation between common measures of assembly quality.** Significant correlations between measures of sequence correctness (QV), assembly correctness (estimated number of misassemblies per kilobase (kb)), assembly completeness (Merqury's Completeness and BUSCO score (including partial and total BUSCOs)), and assembly contiguity (e-size, and N50 )(Spearman's ρ; $p < 0.05$).

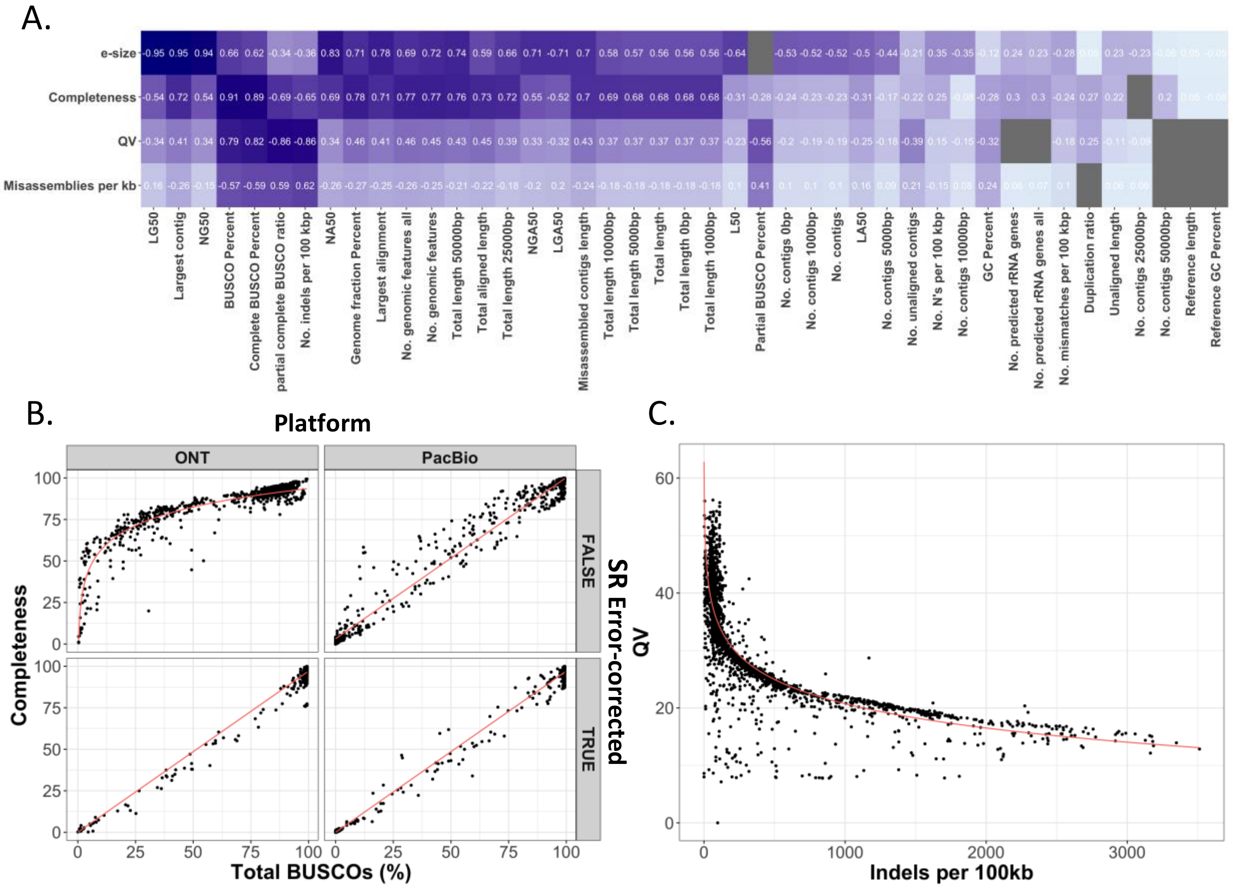


**Figure S2. Correlation between short-read- dependent and independent assembly metrics.** A) Significant correlations between main short-read-dependent estimates of assembly quality used in this study and short-read-independent assembly metrics generated using Quast (Spearman's ρ; $p < 0.05$). Grey boxes represent non-significant correlations. B) ONT-reads not short-read (SR) error-corrected (upper-left) generate assemblies with a fundamentally different relationship between total fraction of identified BUSCOs (partial plus complete or "BUSCO Percent") and Merqury's completeness score (Red lines = Fitted linear models; Upper-left: $y = \alpha + ßlog(x)$, adj. $R^2 = 0.86$, $p < 2.2e\text{-}16$; Other panels: $y = \alpha + ßx$, adj. $R^2 = 0.98$, $p < 2.2e\text{-}16$) C) Correlation between BUSCO identified indels and Merqury's QV (Red line = fitted linear model; $y = \alpha + ßlog(x)$, adj. $R^2 = 0.66$, $p < 2.2e\text{-}16$).

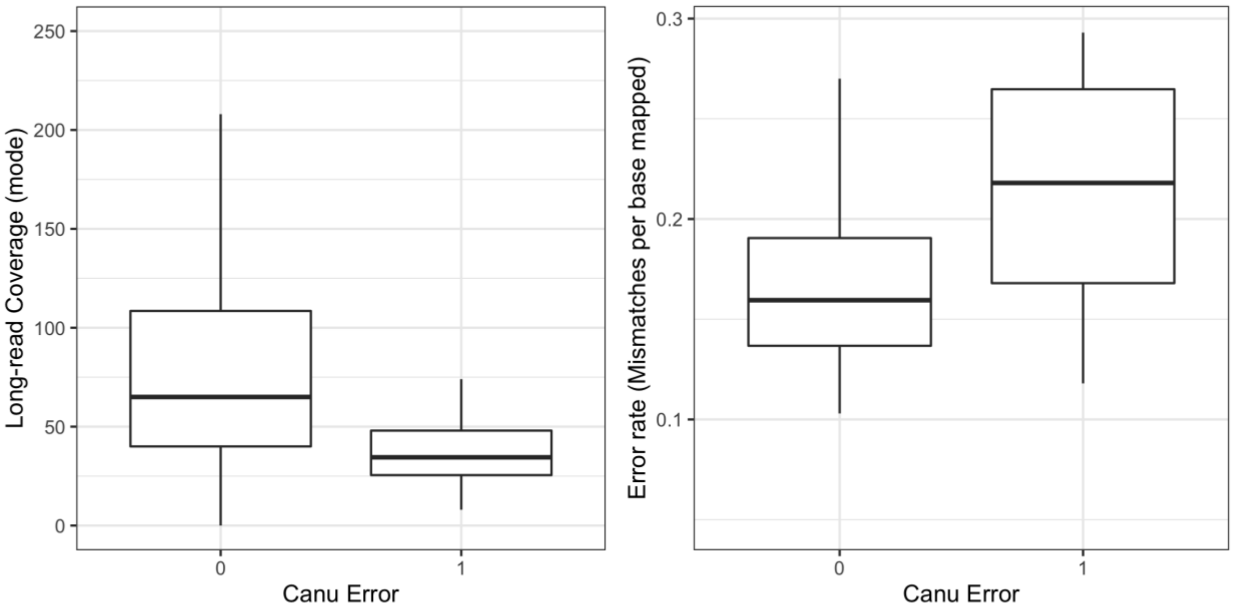


**Figure S3. Canu assembly errors (program failures) are associated with low coverage and higher error rates in raw reads.** Distribution of long read coverage modes (a) and error rates (b) for data sets from which Canu was able to generate assemblies regardless of error correction method (0) and for data sets that generated Canu errors in at least one instance (1).

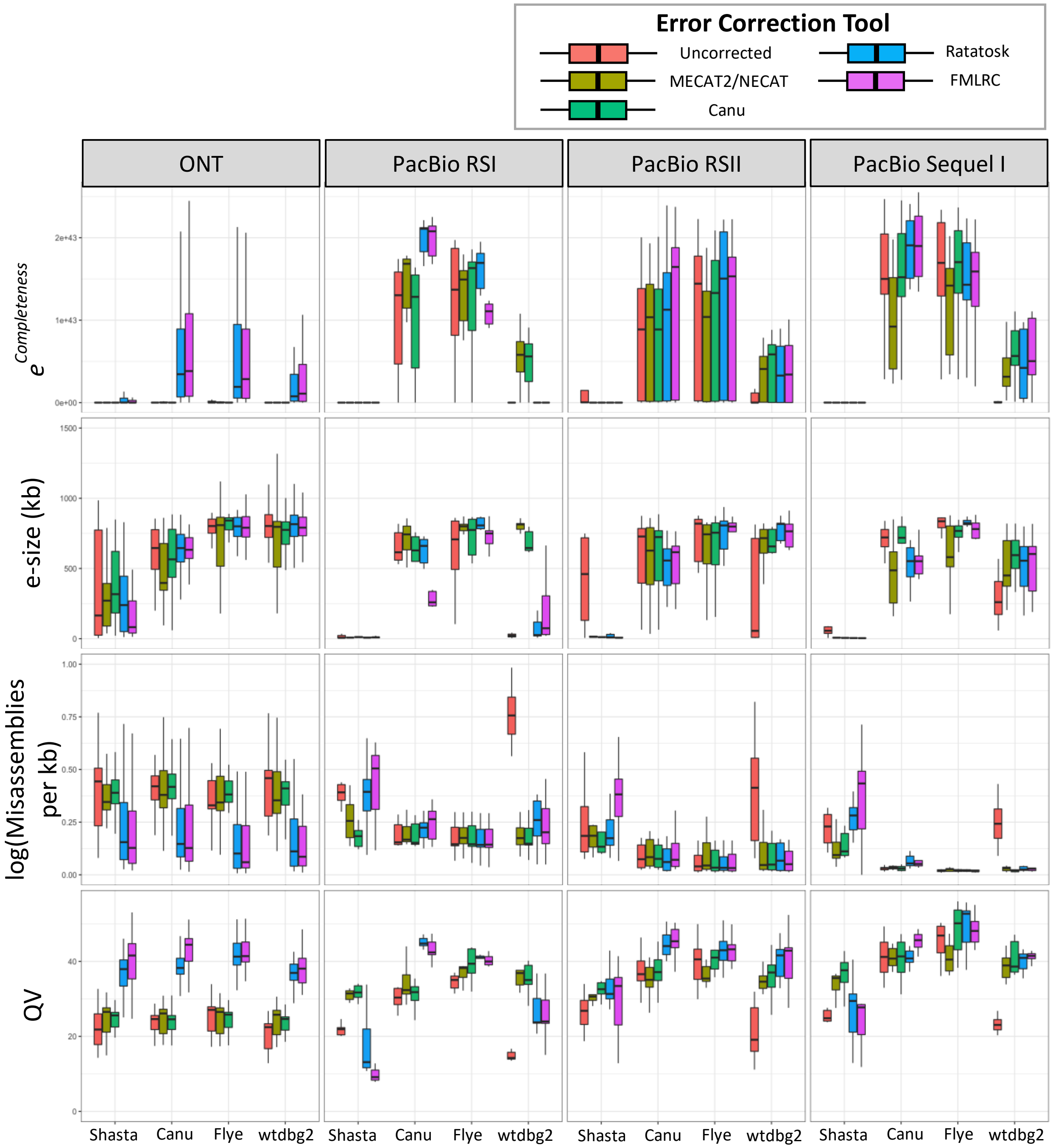


**Figure S4. Distributions of assembly quality metrics based on error correction tool and assembler program.**

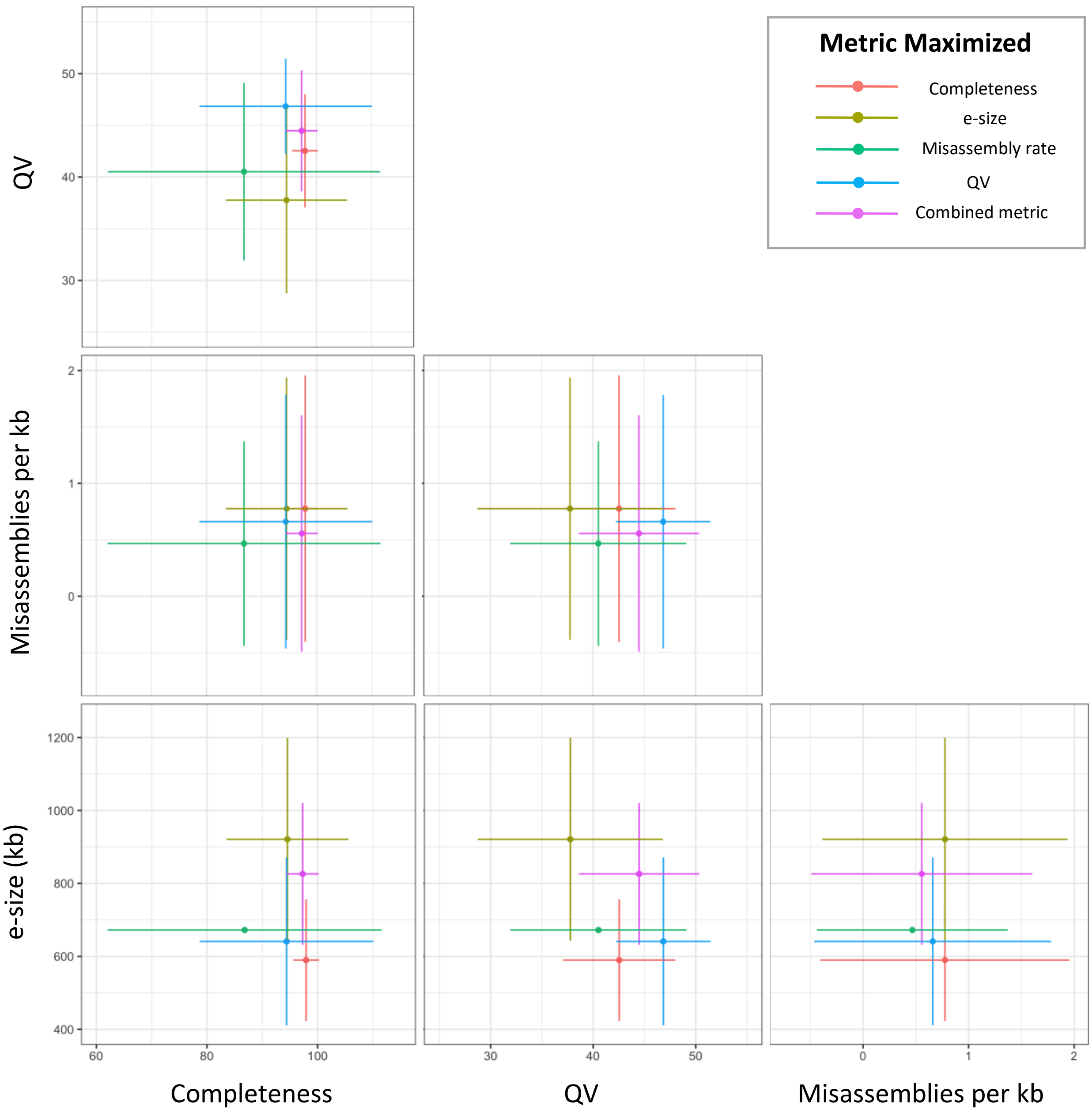


**Figure S5. Tradeoffs between assemblies optimized for each assembly quality metric.** Each point shows the mean and standard deviation of quality values (x and y axes) for assemblies that scored best in terms of completeness, e-size, misassembly rate (green), QV, and across all metrics using unweighted sums of z-scores.

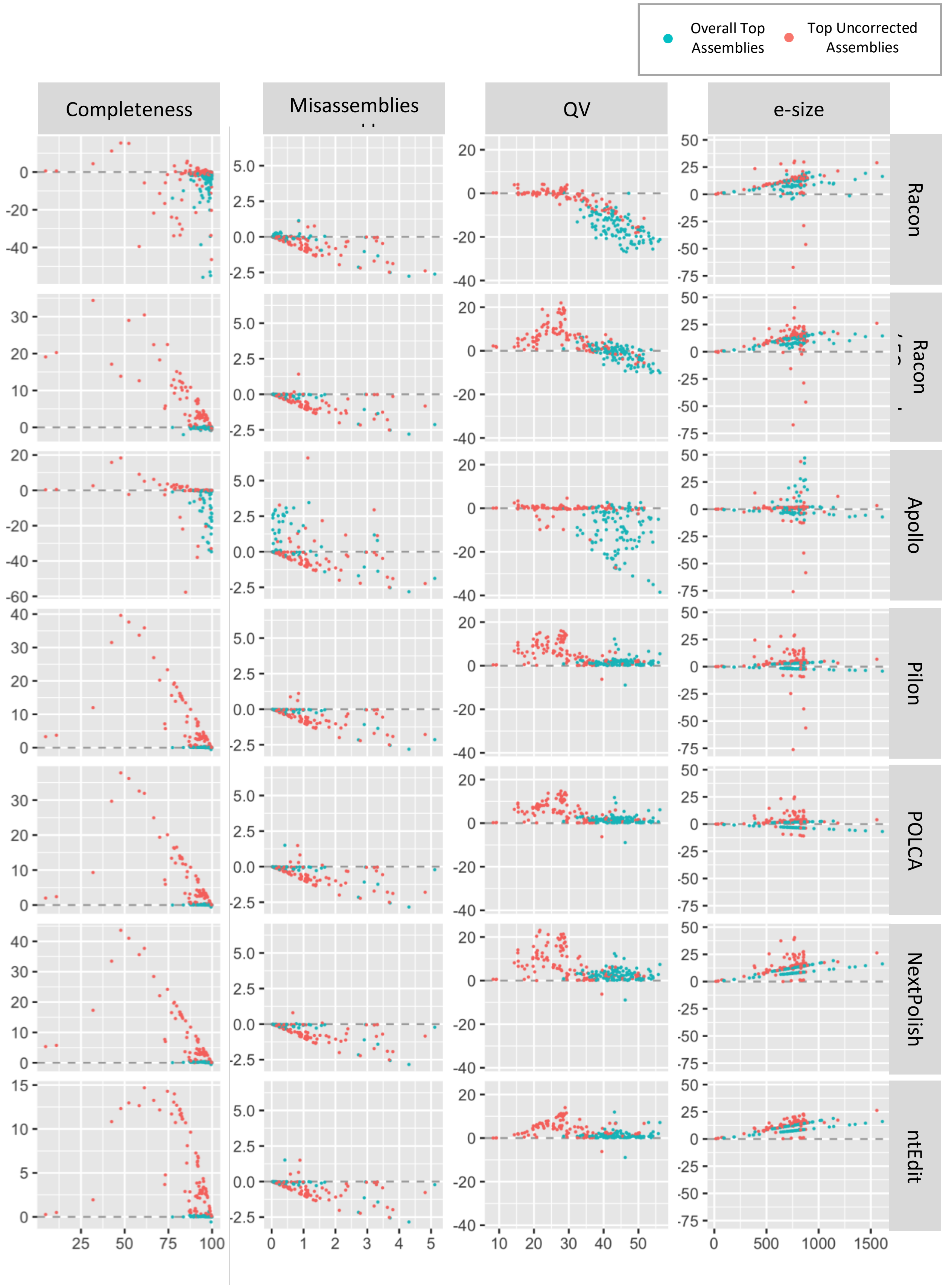

Overall Top Assemblies
Top Uncorrected Assemblies
Completeness
Misassemblies
QV
e-size
Racon
Apollo
Pilon
POLCA
NextPolish
ntEdit

**Figure S6. Correlations between pre-polished assembly quality (x-axis) and changes due to polishing (y-axis) do not follow consistent linear patterns across polishing tools or assembly metrics.** Note that outliers, in terms of Δe-size, greater than 50kb have been removed for the sake of clarity

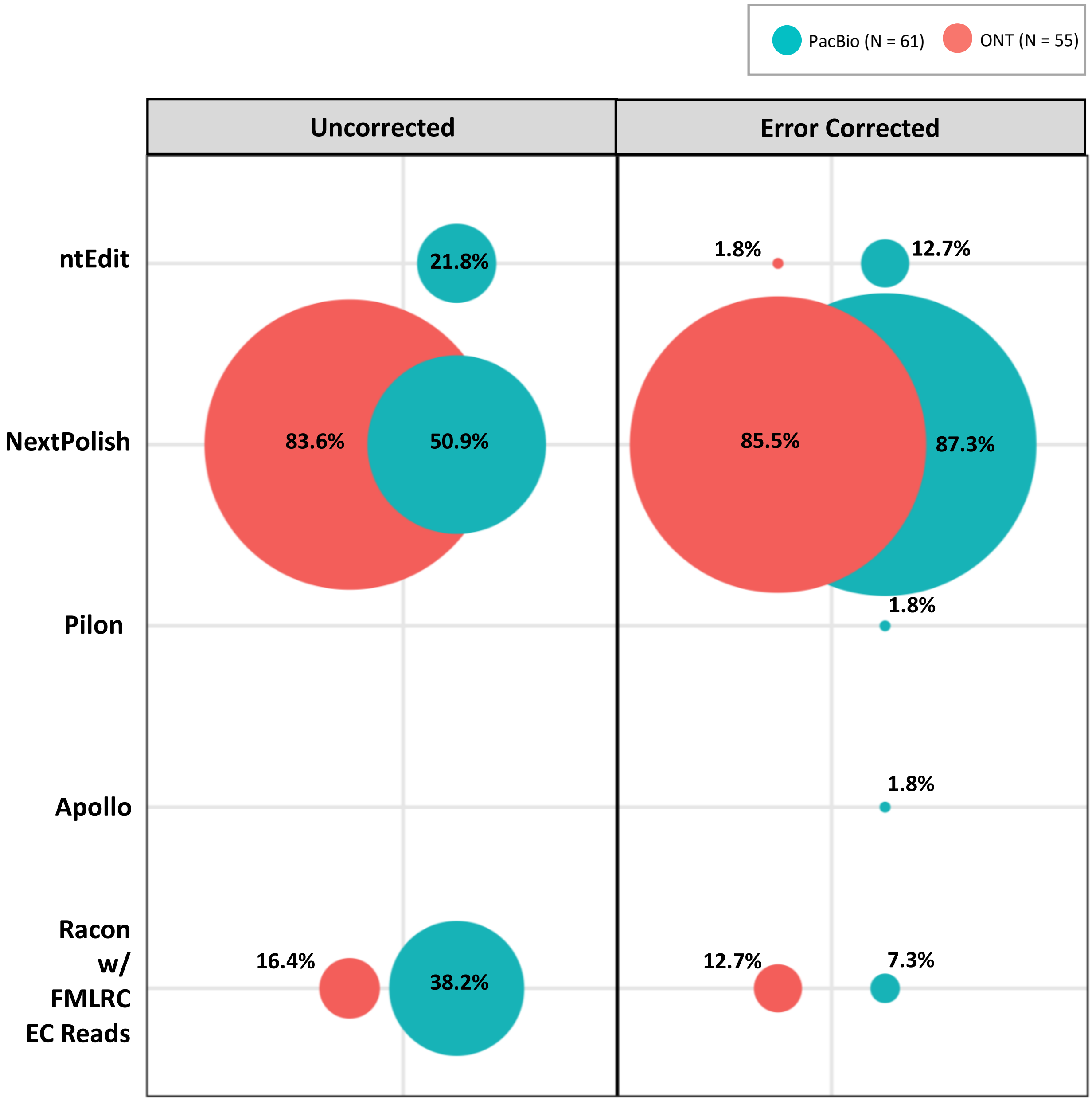


**Figure S7. NextPolish produces the greatest gains in quality among polishing tools in the largest proportion of datasets.** Fraction (corresponding to circle area) of assemblies with the greatest quality gains generated by each polishing tool for assemblies generated with error-corrected and uncorrected input assemblies. Ranking was based on e-size, Merqury completeness, and Merqury QV.

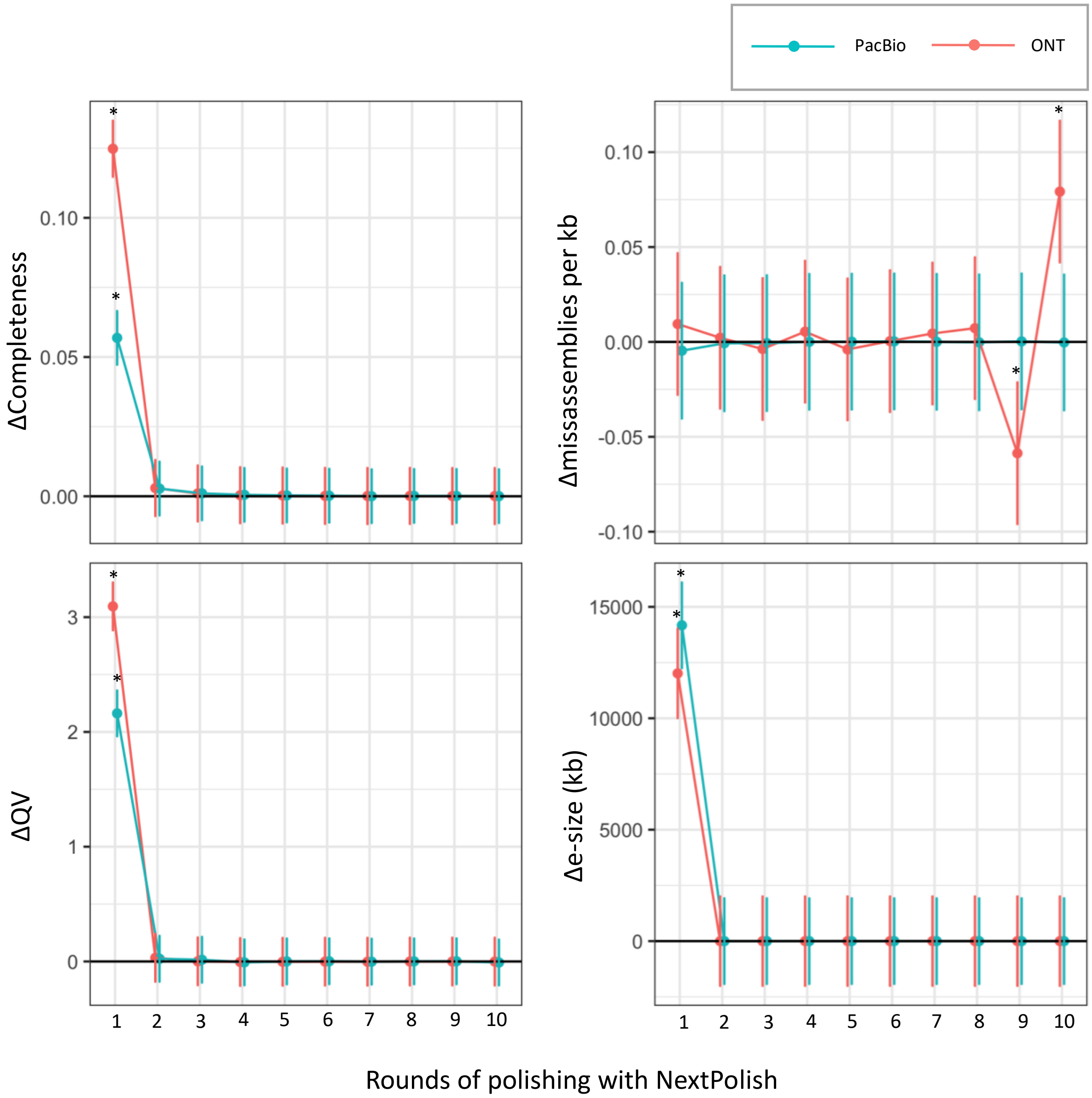


**Figure S8. Effects of multiple rounds of short-read polishing on assembly quality.** Estimated marginal mean change in values of four measures of assembly quality relative to assembly from previous round. Error bars = 95% CI. * = Value is significantly greater or less than 0 (t-test; Bonferroni adjusted $p < 0.05$). Tests based on estimated marginal means derived from multivariate model.

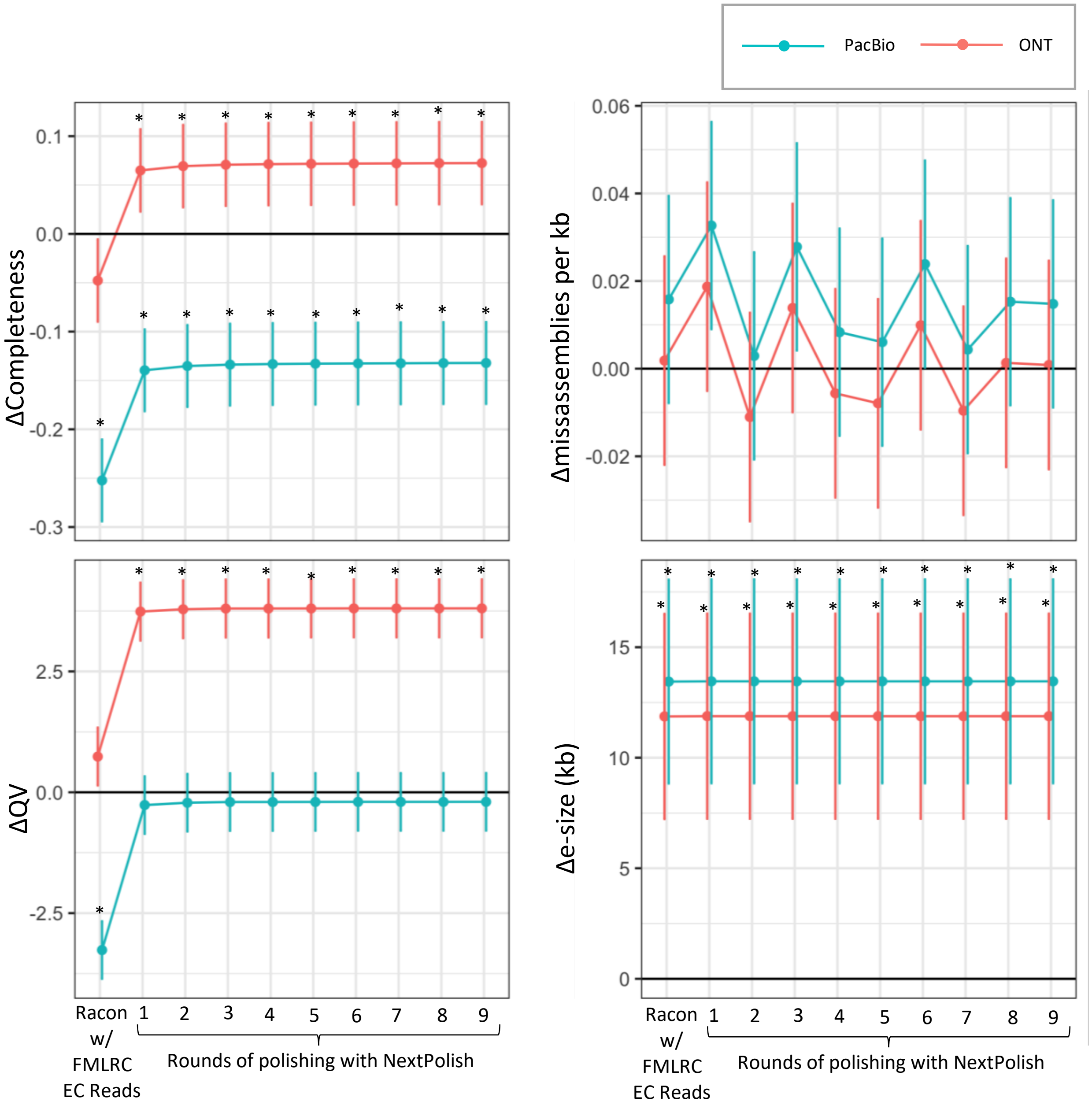


**Figure S9. Effects of combined long-read and short-read polishing.** Estimated marginal mean change in values of four measures of assembly quality *relative to the unpolished assembly*. Error bars = 95% CI. * = Value is significantly greater or less than 0 (t-test; Bonferroni adjusted $p < 0.05$). Tests based on estimated marginal means derived from multivariate model.

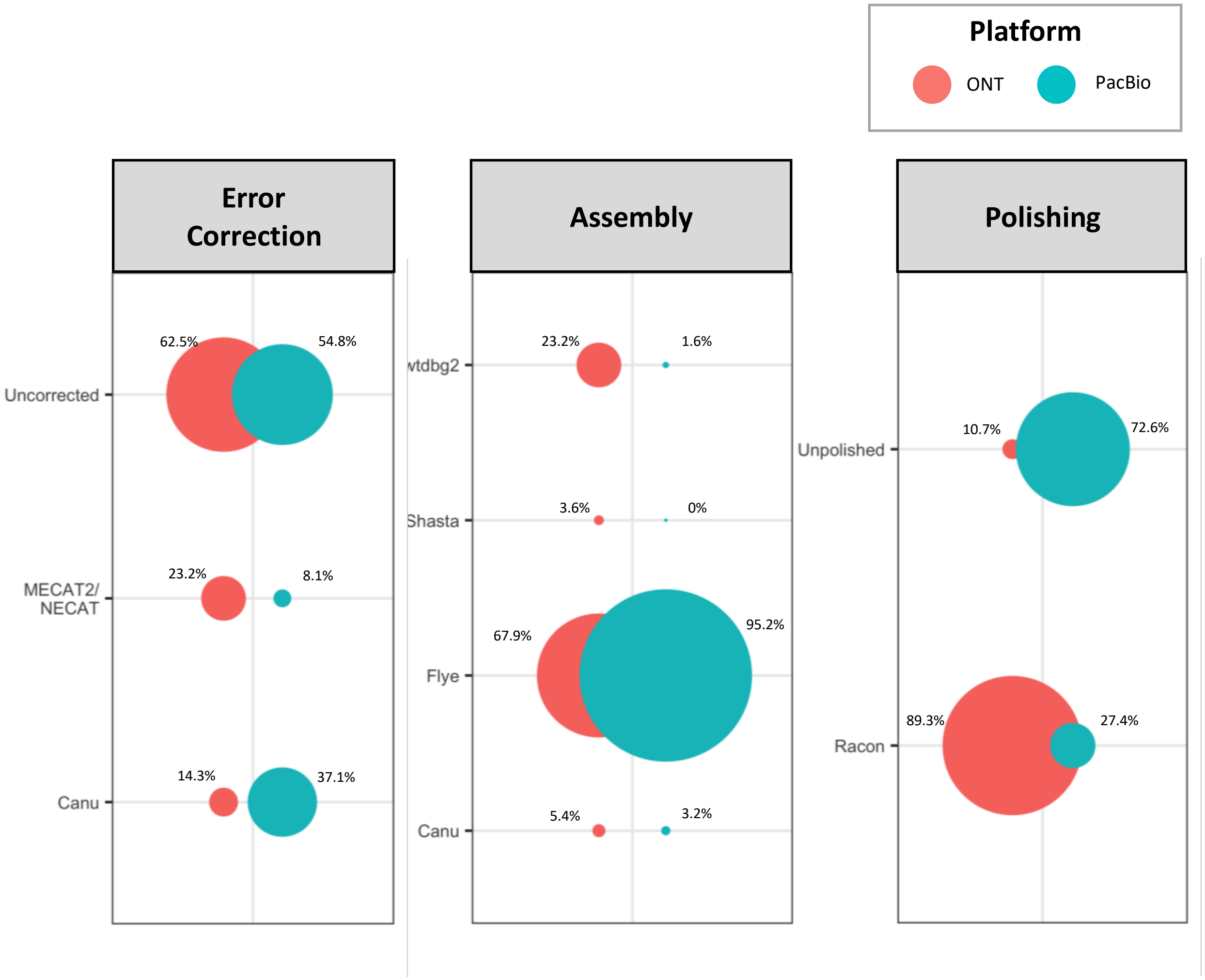


**Figure S10. Top assembly pipelines for assemblies generated without supplemental short reads.** Fraction (corresponding to circle area) of assemblies ranked highest for each error correction, assembly, and polishing tool combination.

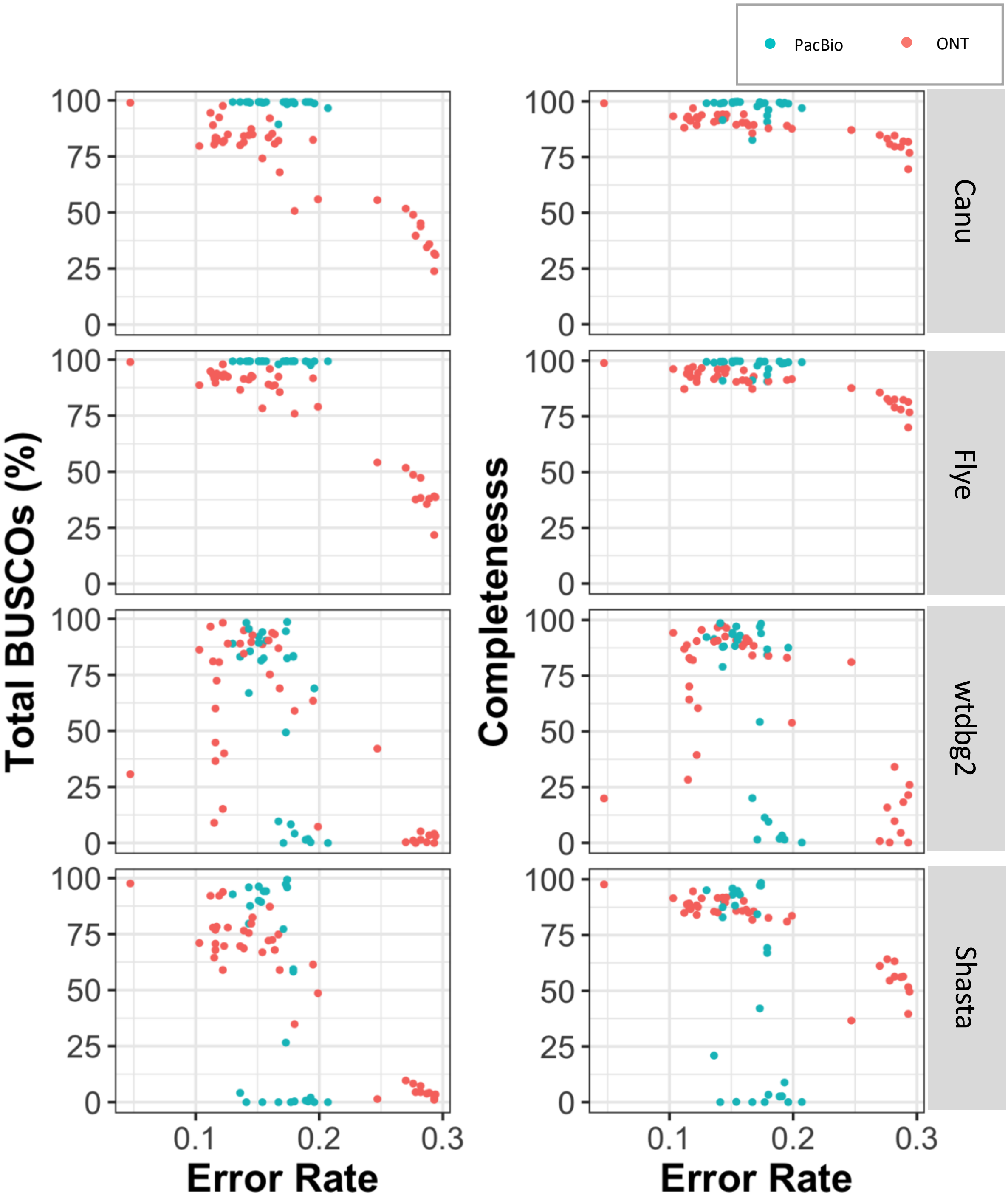


**Figure S11. The percentage of identified BUSCOs is more sensitive to higher error rate data sets than Merqury completeness.** Correlation between long read error rate and completeness metrics for assemblies generated from uncorrected reads.

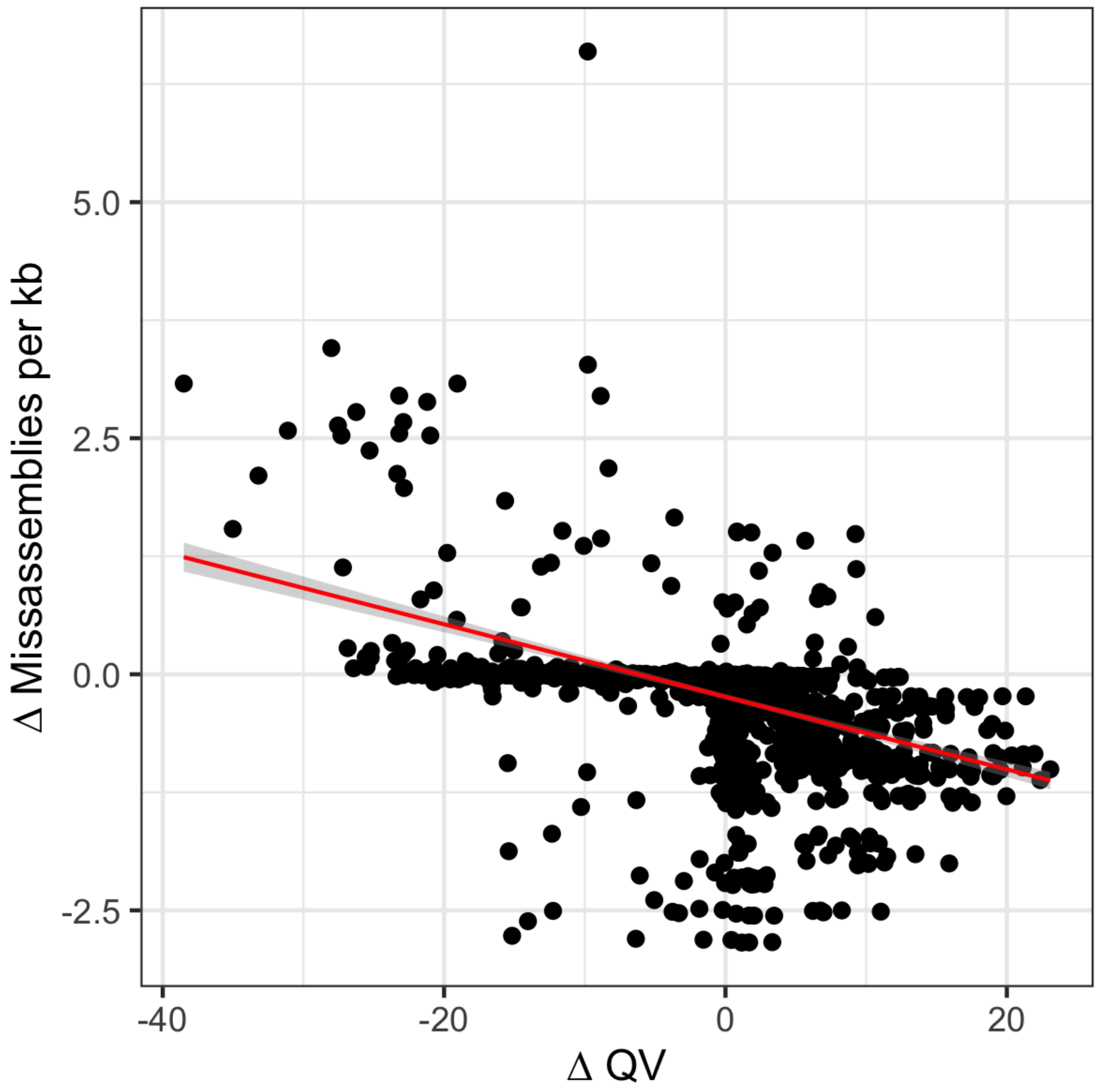


**Figure S12. Polishing effects on QV are correlated with polishing effects on missassembly rates.** Linear model; Adjusted $R^2 = 0.19$; $p < 2.2e-16$.
